# Does over-reliance on auditory feedback cause disfluency? An fMRI study of induced fluency in people who stutter

**DOI:** 10.1101/2020.11.18.378265

**Authors:** Sophie Meekings, Kyle Jasmin, Cesar Lima, Sophie Scott

## Abstract

This study tested the idea that stuttering is caused by over-reliance on auditory feedback. The theory is motivated by the observation that many fluency-inducing situations, such as synchronised speech and masked speech, alter or obscure the talker’s feedback. Typical speakers show ‘speaking-induced suppression’ of neural activation in superior temporal gyrus (STG) during self-produced vocalisation, compared to listening to recorded speech. If people who stutter over-attend to auditory feedback, they may lack this suppression response. In a 1.5T fMRI scanner, people who stutter spoke in synchrony with an experimenter, in synchrony with a recording, on their own, in noise, listened to the experimenter speaking and read silently. Behavioural testing outside the scanner demonstrated that synchronising with another talker resulted in a marked increase in fluency regardless of baseline stuttering severity. In the scanner, participants stuttered most when they spoke alone, and least when they synchronised with a live talker. There was no reduction in STG activity in the Speak Alone condition, when participants stuttered most. There was also strong activity in STG in response to the two synchronised speech conditions, when participants stuttered least, suggesting that either stuttering does not result from over-reliance on feedback, or that the STG activation seen here does not reflect speech feedback monitoring. We discuss this result with reference to neural responses seen in the typical population.

## Introduction

Persistent developmental stuttering is a chronic condition that severely disrupts speech production with frequent and involuntary sound prolongations, silent ‘blocks’ and syllable repetitions. For the 1% of adults worldwide who experience it, there is currently no known intervention that will result in permanently fluent speech production. However, temporary periods of fluency are not uncommon, and in many cases it is possible to deliberately induce fluent speech by changing aspects of the talker’s environment— for example, by asking them to speak in loud masking noise (Cherry & Sayers, 1956; Conture & Brayton, 1975; Murray, 1969; Yairi, 1976), or while hearing their voice pitch-shifted or played back to them at a delay (Kalinowski, Armson, Stuart, & Gracco, 1993; Macleod, Kalinowski, Stuart, & Armson, 1995). However, when the masker or altered feedback stops, stuttering resumes.

This, in conjunction with the functional anomalies seen in auditory cortex in neuroimaging studies of people who stutter, has led some researchers to suggest that stuttering is a manifestation of a central auditory processing disorder (Salmelin et al., 1998), or some difficulty with speech monitoring. Max et al. (Max, Guenther, Gracco, Ghosh, & Wallace, 2004) suggested two possible hypotheses, based on Guenther’s (2006) DIVA model of speech production. First, the underlying internal speech models in people who stutter may be poorly specified in some way. Under this hypothesis, stuttering manifests itself during development when children are unable to update their internal speech models appropriately in response to feedback, and may have a problem with accessing or forming mappings between motor commands and sensory responses. As a result their internal model is mis-specified and sends inaccurate feedforward commands to the articulators. The mismatch between the faulty prediction and the actual sensory consequences of the executed movement results in attempts to correct the speech by reissuing the motor command, resulting in stuttering. Altered feedback induces stuttering, therefore, because it activates auditory cortex and stimulates the internal model.

Another model that develops the idea of stuttering as an internal model deficit is the Covert Repair Hypothesis, or CRH (Postma & Kolk, 1993). The CRH uses Levelt’s (1983) three-loop monitoring system as a theoretical frame, rather than Guenther’s DIVA model; however, Levelt’s internal monitoring loop (defined as the inspection of the articulatory plan) and Guenther’s feedforward loop are both conceptually similar in that they describe a stage of speech monitoring that occurs before articulation. The Covert Repair Hypothesis suggests that disfluencies arise because the speaker has detected an error during internal monitoring and is attempting to correct it. Such ‘covert repairs’ occur more frequently in people who stutter owing to a deficit in phonological encoding. This theory is based on the spreading-activation account of phonemic control (Dell, 1986), in which word selection is accomplished by activating all phonemic, semantic and syntactic nodes associated with the word; this activation spreads to surrounding nodes until the most highly activated node is selected. If selection occurs too early, it is more likely that the wrong node will be selected, leading to an error, which in turn triggers a covert repair. Postma & Kolk (1993) argue that PWS are slow to activate the right representation, so are more likely to make these errors. However, evidence does not support the idea that PWS have a phonological disorder: children who stutter make the same amount of phonological errors as fluent children, and the number of phonological errors made does not correlate with stuttering severity (Nippold, 2002). Additionally, evidence suggests that adults who stutter do not have a slower rate of phonological encoding compared to fluent adults (Brocklehurst, 2008).

As an alternative, the second theory put forward by Max et al. (2004) involves no problems with the internal model. Rather, PWS may have weakened feedforward projections and are thus forced to rely on feedback monitoring. Overreliance on feedback monitoring results in system resets and effector oscillations as the talker attempts to compensate for the time delay between the motor command being issued and the feedback being received. Under this hypothesis, altered auditory feedback prevents the talker from relying on the feedback circuit and encourages them to use the weakened feedforward projections, resulting in fewer corrections. A computational modelling study testing this theory using the DIVA model (Civier, Tasko, & Guenther, 2010) found that programming the model to rely more on auditory feedback resulted in more acoustic errors than the default model parameters, particularly during rapid formant transitions. The model did not produce stuttered speech with these parameters; however, the auditory errors produced are consistent with those found in human studies (Blomgren, Robb, & Chen, 1998; Robb & Blomgren, 1997), which found that people who stutter have significantly lower F2 values when producing syllables with rapid formant transitions compared to syllables without rapid transitions. The authors (Civier et al, 2010) suggest that an accumulation of these errors eventually causes a system reset, in which the syllable is restarted, leading to sound and syllable repetition. One interesting observation that may support the theory that stuttering is related to over-reliance on auditory feedback is the existence of several surveys suggesting that there is a much lower incidence of stuttering in the deaf population than in the population at large, although this evidence is largely anecdotal (Backus, 1938; Harms & Malone, 1939; Wingate, 1970).

Here we investigate the hypothesis that results from an over-reliance on auditory feedback (Civier et al., 2010), so choral speech may prevent this by altering the talker’s perception of her own voice. Speech feedback monitoring is hypothesised to take place in the superior temporal gyrus (STG). Typical speakers show ‘speaking-induced suppression’ of neural activation in STG during self-produced vocalisation, compared to listening to recorded speech. If people who stutter over-attend to the sound of their own voice, they may lack this suppression response. We focus on choral speech, in which the person who stutters (PWS) talks in synchrony with another person. This technique reliably causes a dramatic increase in fluency, which is usually greater than that demonstrated in other fluency-enhancing conditions (Johnson & Rosen, 1937; Kiefte & Armson, 2008), and is highly consistent across subjects and studies (Barber, 1939; Ingham et al., 2006; Kalinowski & Saltuklaroglu, 2003). We sought to understand why this is such an effective intervention when other techniques that have been theorised to work in the same way are much more variable.

## Methods

### Participants

Participants were recruited through the British Stammering Association and were adults who self-identified as people who stutter. 21 participants (8 female; mean age 38.7, s.d. 12.2, range 24-63) underwent behavioural pretesting to classify their stuttering severity and evaluate the effect of choral speech on their stutter. Participants were additionally screened for hearing loss using an Amplivox 116 Screening Audiometer with DD45 earphones (amplivox.ltd.uk). One participant met the critera for hearing loss (defined here as a four-frequency pure tone average threshold of more than 20dB) and was excluded from the experiment for this reason.

They were invited back to participate in the fMRI study if they met fMRI safety standards, had a stutter of any severity as defined by the SSI-IV, and they became more fluent under choral speech conditions. One participant was excluded at this stage because they did not stutter during the behavioural test. Of those who were invited back, thirteen native British English speakers continued to the fMRI testing (4 female; mean age 34.7, s.d. 8.4, range 24-48).

### Assessment for stuttering severity

Participants’ speech was evaluated using the Stuttering Severity Instrument IV (Riley, 1972). The SSI-IV calculates a severity score based on the percentage of syllables stuttered in two speech tasks, the duration of the three longest stuttering incidents, and physical tics observed at the time of testing. Participants sat in a soundproofed room with two experimenters. One experimenter delivered the test materials while the second recorded information on physical concomitants. Participants’ speech was recorded using a RODE NT1-A one-inch cardoid condenser microphone connected to a Windows computer via a Fireface UC high-speed USB audio interface (RME Audio, Haimhausen) Their voices were recorded at 44100Hz with 16 bit quantisation using Adobe Audacity 3.0.

Subjects spoke spontaneously for three minutes and read one of two passages aloud. The passages were either 369 or 374 syllables long. The other passage was used in the synchronous speech task and the order of the passages was counterbalanced across participants.

### Synchronous speech outside the scanner

To evaluate the effects of synchronous speech on testing, participants read the second passage in unison with an experimenter positioned outside the testing room. The experimenter spoke into an AKG 190E cardoid dynamic microphone and heard through AKG K240 Studio on-ear headphones. This mimicked the effect of speaking in the scanner environment as the participant was unable to see their conversational partner and use nonverbal cues. It additionally enabled us to record the participant’s voice on its own, without the experimenter.

### FMRI stimuli

The fMRI paradigm was closely based on Jasmin et al. (2016), with some difference in the technical setup and a speech in noise condition substituted for the ‘Diff-Live’ condition.

Participants lay supine in the scanner and saw sentences in yellow or blue on a black background projected onto an in-bore screen, using a video projector (Eiki International). They spoke into an OptoAcoustics FOMRI-III noise-cancelling optical microphone and heard stimuli through Sensimetrics S14 fMRI-compatible insert earphones. In the control room, the experimenter was seated in front of a RODE NT1-A 1” cardoid condenser microphone and heard the participant through Beyerdynamic DT100 circumaural headphones. The participant’s voice, experimenter’s voice and sound from the computer were routed through an RME Fireface UC 36-Channel, 24 Bit / 192 kHz USB high speed audio interface using TotalMix software and were recorded in three separate channels on a Mac computer. Routing was instantaneous, so there was no delay between the experimenter or participant speaking and their conversational partner hearing them.

### FMRI Task

The following five sentences were used as stimuli:

1. When sunlight strikes raindrops in the air, they act as a prism and form a rainbow.
2. There is, according to legend, a boiling pot of gold at one end of a rainbow.
3. Some have accepted the rainbow as a miracle without physical explanation.
4. Aristotle thought that the rainbow was a reflection of the sun’s rays by the rain.
5. Throughout the centuries, people have explained the rainbow in various ways.

These sentences are adapted from The Rainbow Passage (Fairbanks, 1960), and were used as they are about the same length (mean syllables = 20.8 ± 1.3), and can be spoken comfortably during a short presentation window by typical speakers. It was expected that some participants who stuttered would not be able to complete the entire sentence in the six seconds allotted for the task, and this was factored into the analysis.

In every trial, participants saw a prompt telling them which condition was coming up next, followed by the text of one of the five sentences. Instruction prompts were displayed for three seconds, then replaced with a fixation cross which remained on screen for one second before the stimulus sentence was shown. There were six conditions:

1. Synch-Live: Participants saw a ‘SYNCHRONIZE’ prompt and read the sentence synchronously with the experimenter.
2. Synch-Rec: Participants saw a ‘SYNCHRONIZE’ prompt and read the sentence synchronously with a recording of the experimenter.
3. Speak-Noise: Participants saw a ‘SPEAK IN NOISE’ prompt and read the sentence over 83dB white noise.
4. Speak-Alone: Participants saw the prompt, ‘SPEAK’ and read the sentence on their own
5. Listen: Participants saw the prompt, ‘LISTEN’ and read the sentence silently while hearing a recording of the experimenter reading it aloud.
6. Read-Silently: Participants saw the prompt ‘READ SILENTLY’ and read the sentence silently with no auditory stimulus.

In the synchronization conditions, participants spoke with a male American English speaker, either live, through the microphone (Synch-Live) or recorded, via a laptop (Synch-Rec). Recorded trials in both Synch-Rec and Listen conditions were produced by the live experimenter during synchronous speech with a different partner. This was intended to isolate neural and behavioural correlates of speech with a live partner who can adaptively alter their voice to match yours (reciprocal synchronization) while controlling for auditory and motor requirements as closely as possible. The prompt for both synchronization conditions was identical apart from a colour code intended to tell the experimenter when live speech was required: the prompt text was yellow in the Synch-Live condition, and blue in the Synch-Rec condition. To disguise the colour code from participants, the colour of the prompts was varied randomly in all other conditions, so that the prompt was blue in half of all trials, and yellow in the rest. Post- test debriefing confirmed that none of the participants identified that they were synchronizing with a recording.

### FMRI acquisition parameters

Functional MRI images were acquired using a Siemens Avanto 1.5 Tesla scanner with 32-channel head coil, using a T2-weighted gradient-echo planar imaging sequence, which covered the whole brain (TR=9s, TA=3s, flip angle 90 degrees, 35 axial slices, matrix size=64×64×35, 3×3×3mm in-plane resolution). High-resolution anatomical volume images (HIRes MP-RAGE, 160 sagittal slices, matrix size: 224×256×160, voxel size=1 mm^3^) were also acquired for each subject. Participants took part in three functional runs, each consisting of 55 trials (ten of each main condition and five ReadSilently trials). The five stimulus sentences were crossed with each of the six conditions, and combination of stimulus sentence and condition appeared twice per run. The order of the trial types was pseudo-randomized such that every five trials included one of each of the five stimulus sentences and one trial in each condition.

## Acoustic and behavioural analysis

### Behavioural pretesting

Participants’ speech was transcribed and rated for number of stuttered syllables, perceived naturalness (on a scale from 1-9, with 9 being ‘highly unnatural’) and the duration of the longest stuttered syllable. FDR-corrected paired t-tests were used to evaluate the difference between solo reading and choral reading for each parameter.

### FMRI recordings

Two participants’ recordings could not be used owing to problems with the recording setup. For the remaining participants, recordings of each experiment were divided up into individual trials using a MATLAB script. Each trial was evaluated for the total number of syllables, and the number of stuttering events, by a rater who was blind to the conditions. These scores were used to generate average speech rates (in syllables per second) and percentage of stuttered syllables for each participant in each condition. A within-subjects ANOVA was carried out to evaluate differences in dysfluency rate between conditions.

## Functional analysis

### Preprocessing

First-level and group-level analysis was carried out using SPM 8. To allow for T1 saturation effects, the first three functional volumes of each run were discarded. Each participant’s fMRI time series was realigned to the first volume of the run using six-parameter rigid-body spatial transformation and their mean functional image was coregistered to their anatomical T1 image; the scans were then re-oriented into standard space by manually aligning to the anterior commissure. The estimated parameters resulting from motion correction were inspected and did not exceed 3mm or 3 degrees in any direction. The T1 image was segmented into grey matter, white matter and cerebrospinal fluid; the parameters generated by this were used to spatially normalize the functional images into MNI space at 2mm^3^ isotropic voxels. The data was then smoothed using a Gaussian kernel of 8 mm^3^ FWHM.

### Univariate functional analysis

At the single-subject level, events were modelled from the presentation of the stimulus sentence, using a canonical haemodynamic response function, with ReadSilently as an implicit baseline and motion parameters included as a regressor of no interest. Event duration was set at six seconds. Contrast images were calculated for each of the conditions using ReadSilently as a baseline, and for Synch-Live>Synch-Rec.

These contrasts were taken up to the group level and used to perform 1) a one-sample t-test for SynchLive>Synch-Rec; 2) a one-way repeated measures ANOVA looking at differences between each of the three speaking tasks (SpeakAlone, Synchronize, and SpeakNoise) compared to listening. Next, a multiple regression analysis was carried out on the subset of subjects for whom audio data was available (7 subjects) using the percentage of stuttered syllables in each trial as a regressor; this analysis revealed voxels that were more active when participant stuttered, regardless of the trial type. All contrasts were thresholded using a voxel wise familywise error rate correction for multiple comparisons at p <0.05. Statistical images were rendered on the normalized mean functional image for the group of participants.

## Results

### Behavioural pretesting

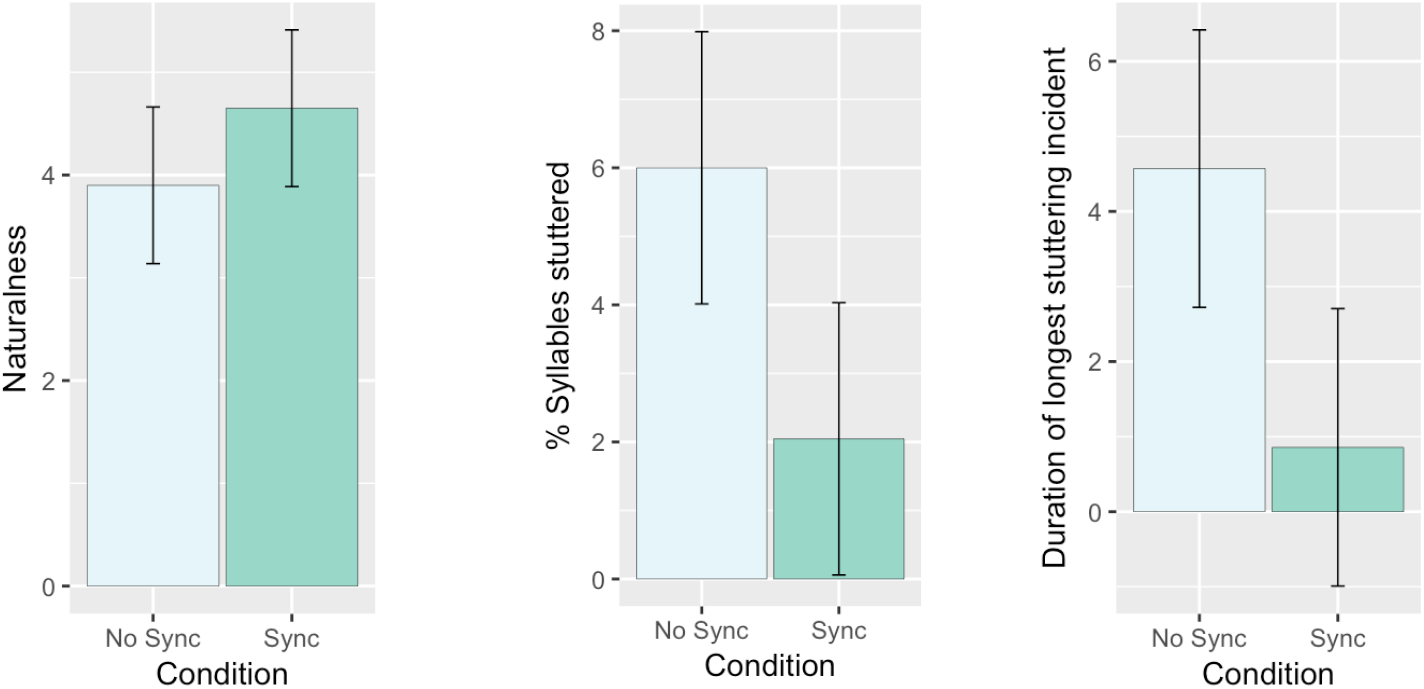

Participants were classified according to the SSI-IV using the recordings made during behavioural pre-testing, and represented a broad spectrum of stuttering severity from very mild to very severe.

A series of one-tailed t-tests were conducted to investigate differences in stuttering duration, frequency and speech naturalness. These revealed that when synchronizing, participants stuttered less than when speaking alone (t(19)=2.149, p=0.025, d=0.51), and the duration of the longest stuttering incident was significantly shorter when synchronising (one-tailed t(19)= 1.987, p=0.034, d=0.48). However, there were no significant changes in speech naturalness between conditions (t(19)=0.099, p=0.46, d=0.02). In the speak-alone condition there was considerable variability in the percentage of stuttered syllables (mean= 7.5, s.d.=11.39) and duration of stuttering incidents (mean= 4.87, s.d.=7.48). By contrast, there was relatively little variability in participants’ performance during choral speech, either in percent stuttered (mean=0.93, s.d.=0.73) or in duration (mean=0.83, s.d.=0.53)

### fMRI data

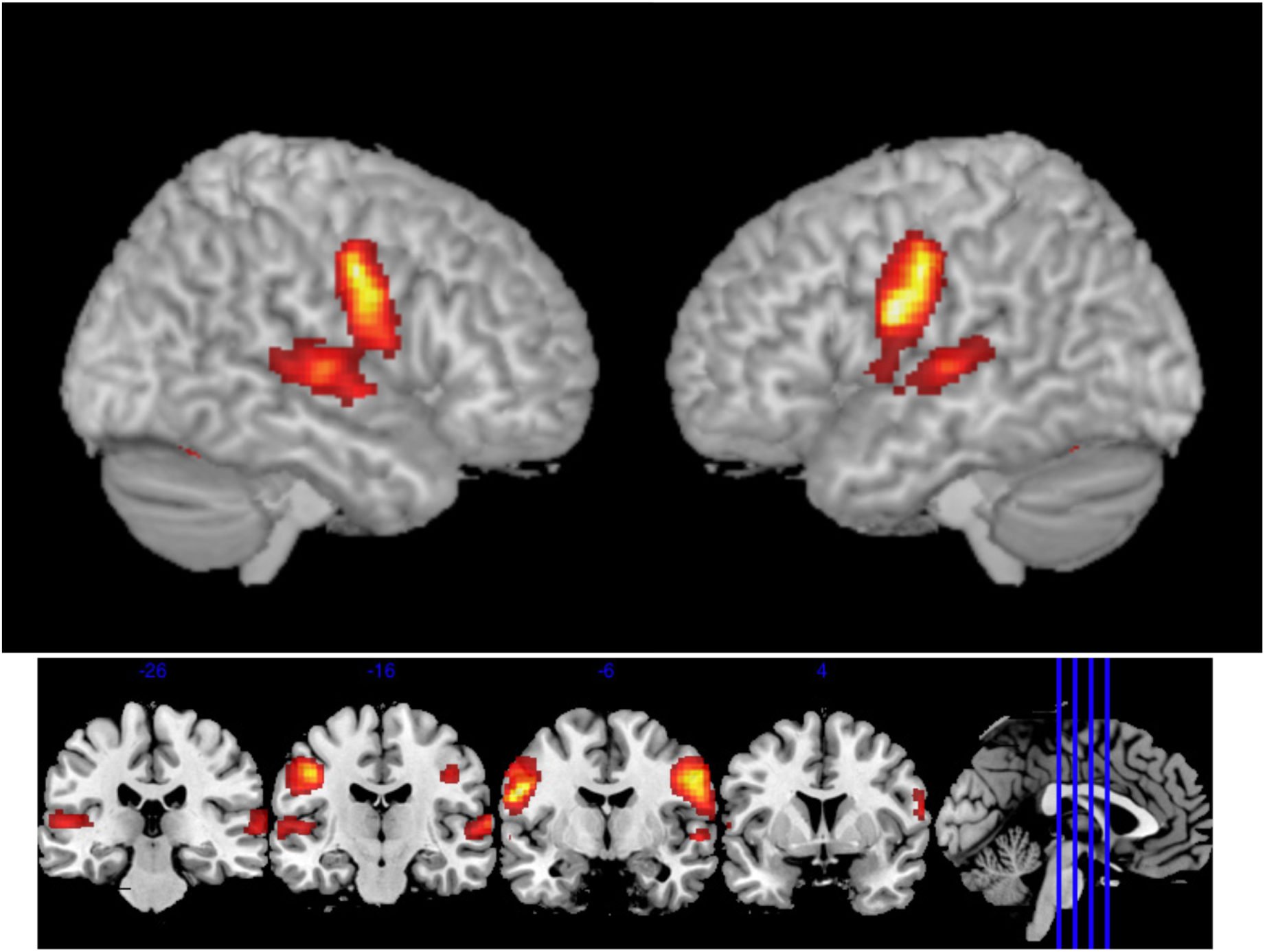

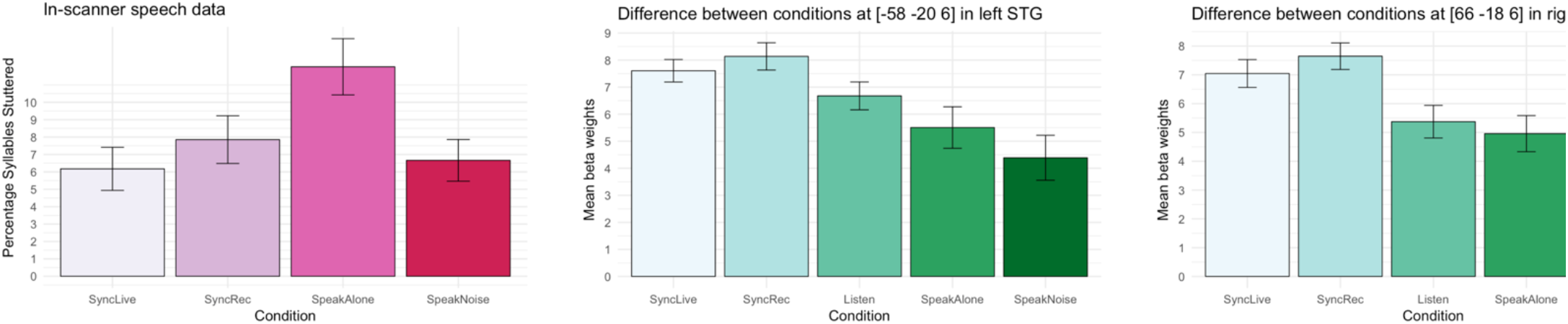

An ANOVA examining areas of the brain where there were significant differences between one or more of the experimental conditions (Listen, SpeakAlone, SpeakNoise and Synchronize, with ReadSilently as an implicit baseline) revealed widespread activation in bilateral superior temporal cortices extending to postcentral gyri, and cerebellum including bilateral Lobule VI and cerebellar vermis. Additional, smaller clusters were seen in the basal ganglia including thalamus, and parietal, occipital and frontal cortex. To investigate differences between conditions, mean beta values were extracted from selected peak voxels and analysed in SPSS (IBM). As this involved running multiple tests, p-values were FDR corrected for multiple comparisons using the method described by Benjamini and Hochberg (1995), and are reported as corrected q-values.

Peak beta values at [−58 −18 10] in the left STG and [60 −22 0] in the right hemisphere were compared using a repeated measures ANOVA with Hemisphere and Condition as factors. Assumptions of sphericity were met for the Hemisphere factor (as it has only two levels) and for Condition (non-significant Mauchly’s W, χ2 (5)=3.14, p=0.68), but were violated for the interaction between Condition and Hemisphere (significant Mauchly’s W, χ2 (5)=14.31, p= 0.015), so the Greenhouse-Geisser correction for degrees of freedom was applied (ε=0.49). The F-test revealed a main effect of Condition (F(3,24)= 11.5, q<0.001, ηp2=0.59) and of Hemisphere (F(1,8)= 30.8, q=0.007, ηp2=0.79) but no significant Condition*Hemisphere interaction (F(1.46,11.66)=2.55, q=0.84). Sidak-corrected posthoc t-tests investigating the main effects of Condition and Hemisphere showed that, bilaterally, responses were significantly greater in the Synchrony condition than the during the other three tasks (p>0.05), with no other significant differences between conditions. Responses in this region were significantly greater in the left than the right hemisphere (p= 0.001).

A second ANOVA looked at effects of Condition and Hemisphere in bilateral postcentral gyri at [−46 −12 36] and [54 −8 38]. Mauchly’s test showed that the assumption of sphericity was violated for the main effect of Condition (χ2 (5)=34.1, p<0.001) and the interaction between Condition and Hemisphere (χ2 (5)= 13.7, p=0.019), so the Greenhouse-Geisser correction was applied. There was a significant main effect of Condition (F(1.19, 9.5)= 66.6, q<0.001) but no effect of Hemisphere (F(1,8)= 2.25, q=1.204) or a significant Condition*Hemisphere interaction (F(1.38, 11.06)=6.05, q=0.168). Sidak-corrected post-hoc t-tests found that the main effect of Condition was attributable to significantly greater BOLD responses in the three speaking conditions (SpeakNoise, SpeakAlone and Synchronize) than during Listen. There were no other significant differences between conditions.

Two one-way repeated measures ANOVAs investigated neural responses to Condition in the cerebellum. The first looked at responses in the left cerebellum at peak [−14 −64 −22]. The data did not meet the assumption of sphericity (Mauchly’s W, χ2 (5)=22.5, p<0.001) so the Greenhouse-Geisser estimates of degrees of freedom were used (ε=0.41). The F-test showed a significant main effect of Condition (F(1.22,9.78)=32.43, q<0.001, ηp2=0.80). Sidak-corrected post-hoc tests showed that activation in the Listen condition was significantly lower than in all other conditions (p<0.004). The second F-test investigated the effect of Condition in the cerebellar vermis at peak [−2 −44 −20]. Mauchly’s test was significant, indicating non-sphericity (χ2 (5)=12.59, p=0.029) so the Greenhouse-Geisser correction was applied (ε=0.51). There was a significant effect of Condition (F(1.5,12.1)=7.11, q=0.013, ηp2=0.47), which post-hoc Sidak corrected t-tests demonstrated was owing to a significantly lower response in the Listen condition than in SpeakNoise (p=0.028).

## Discussion

Behavioural performance both in and out of the scanner confirmed that synchronised speech is extremely effective at inducing fluency in people who stutter, regardless of stuttering severity. Subjects’ speech contained fewer stuttering incidents, and the incidents were of shorter duration, when they spoke chorally compared to speaking alone. Masking noise also reduced the percentage of syllables stuttered and, contrary to expectations, there were no significant differences in measures of stuttering severity when participants spoke in noise compared to when they synchronised with a partner, suggesting that in this experiment, both altered feedback techniques were equally effective at inducing fluency. The two synchronous speech conditions, SyncLive and SyncRec, had similar effects on fluency, but analysis of the recordings showed that participants synchronized more effectively when they spoke with a live experimenter than when they were synchronizing with a recording, confirming the effect identified by Jasmin et al (2016). However, the functional analysis failed to find a neural distinction between the two synchrony conditions. Jasmin et al (2016) found that synchronising with a recording was associated with suppression in temporal cortex relative to listening to sounds, while synchronising with a live partner resulted in a release from this suppression. Here, a univariate analysis showed that responses in the STG bilaterally were the same for speaking alone, listening to speech, and speaking in masking noise, but significantly greater when participants spoke in synchrony either with a live partner or with a recording. Note however that we did not directly replicate Jasmin et al’s analysis (a region of interest analysis limited to the right temporal pole), potentially explaining the difference in results. Our results suggest that PWS do not display a speaking-induced suppression response, which may support the theory that stuttering arises from an over-reliance on auditory feedback. However, if this is the case and disfluency is related to STG over-activation then it is unclear why synchronous speech, which induces fluency, should be associated with an increase in STG activation. It is possible that the response to choral speech in the STG could reflect a preferential response to informational masking, such as that found in Meekings et al., 2016. This is especially plausible as, to synchronize effectively, it was necessary for participants to attend closely to the speech of their conversational partner.

We also saw activation in the cerebellum associated with the different speech production conditions. Further analysis demonstrated that activity in several cerebellar regions was positively correlated with stuttering severity. This included the cerebellar vermis, which has previously been implicated in dysfluency (Brown, Ingham, Ingham, Laird, & Fox, 2005), supporting Budde, Barron & Fox’s (2014) finding that activation in cerebellar vermis is associated with state stuttering (though c.f. (Belyk, Kraft, & Brown, 2015). A possible future analysis integrating neural and behavioural data could look at activation associated with natural fluency (that is, fluency in the SpeakAlone condition) versus activation associated with induced fluency (fluency in the SpeakNoise and choral speech conditions), as suggested by Budde et al. (2014).

This experiment was designed to test the theory that stuttering results from an over-reliance on auditory feedback. The evidence on this point is inconclusive. On the one hand, we found over-activation in the STG when participants spoke alone, which appears to support the over-activation hypothesis. However, synchronised speech was associated with an even greater STG response, which is unexpected if the STG supports error monitoring. Based on our previous finding that activation in the STG is modulated by informational masking during speech production (Meekings et al., 2016), it is likely that we are seeing evidence for multiple streams of processing in the STG.

The results of our other analyses additionally point to a role for the basal ganglia and cerebellum in stuttering, consistent with previous research (Belyk et al., 2015; Giraud et al., 2008). Activity in these areas was associated with speaking alone and talking over a noise masker, and was correlated with stuttering severity. Since these regions are involved in the timing of self-paced movement, this may provide evidence that stuttering arises from a deficit in movement timing and regulation. However, it should be noted that speech rate did not significantly differ between altered speech conditions, and was significantly slower when participants spoke alone (and stuttered), rather than being positively associated with increased fluency as might be expected.

There were a number of differences between this study and previous research, most notably our finding that PWS did not display a speaking-induced suppression response-that is, the STG was over-active when participants spoke alone, rather than under-active as previously suggested (Brown et al, 2015). Additionally, despite previous studies suggesting that activity in auditory and motor cortex is right-lateralized in PWS, in our sample we found that peak activity was greater in the left hemisphere than in the right. To confirm these results, it would be desirable to recruit a larger sample of people who stutter, as well as a control group-unfortunately, constraints on time and resources meant that this was not possible at the time of testing. It should be noted that there is disagreement even among large-scale meta-analyses about the neural hallmarks of stuttering as a trait or state (Budde et al., 2014; Belyk et al., 2014). Stuttering may be an ‘umbrella syndrome’ composed of multiple disorders with overlapping symptoms but distinct aetiologies. For example, stuttering can be caused by head injury (Alm, 2004) and there is considerable individual variability within subjects (Wymbs, Ingham, Ingham, Paolini, & Grafton, 2013). In this study we found several unique results. In our sample of people who stutter, synchronized speech recruits a distinct network of cortical and cerebellar regions that are not modulated by other types of speech production. Additionally, our analysis showed that the STG does not distinguish between hearing sounds, speaking alone, and talking in masking noise, lacking the speaking-induced suppression response seen in typical speakers. However, it responds significantly more to speaking synchronously, potentially reflecting different streams of processing within auditory cortex. Although these results do not provide unequivocal support for the feedback over-reliance hypothesis, they have implications for our understanding of how the STG works and how PWS process speech. Further research with an expanded sample size and control group can confirm our findings and contribute to our understanding of how PWS differ from typical speakers.

